# Postnatal inflammation in *Apoe*^−/−^ mice is associated with immune training and atherosclerosis

**DOI:** 10.1101/2021.04.14.439403

**Authors:** Ellesandra C. Noye, Siroon Bekkering, Albert P. Limawan, Maria U. Nguyen, Lisa K. Widiasmoko, Hui Lu, Salvatore Pepe, Michael M. Cheung, Trevelyan R. Menheniott, Megan J Wallace, Timothy J. Moss, David P. Burgner, Kirsty R. Short

## Abstract

**Background and aims:** Preterm birth is associated with increased risk of cardiovascular disease (CVD). This may reflect a legacy of inflammatory exposures such as chorioamnionitis, that complicate pregnancies delivering preterm, or recurrent early-life infections, common in preterm infants. We previously reported that experimental chorioamnionitis followed by postnatal inflammation has additive and deleterious effects on atherosclerosis in *Apoe*^−/−^ mice. Here, we aimed to investigate whether innate immune training is a contributory inflammatory mechanism in this murine model of atherosclerosis.

**Methods:** Bone marrow-derived macrophages and peritoneal macrophages were isolated from 13 week-old *Apoe*^−/−^ mice, previously exposed to prenatal intra-amniotic (experimental choriomanionitis) and/or repeated postnatal (peritoneal) lipopolysaccharide (LPS). Innate immune responses were assessed by cytokine responses following 24-hour *ex vivo* stimulation with toll-like receptor (TLR) agonists (LPS, Pam3Cys), and RPMI for 24 hours. Bone marrow progenitor populations were studied using flow cytometric analysis.

**Results:** Following postnatal LPS exposure, bone marrow-derived macrophages and peritoneal macrophages produced more pro-inflammatory cytokines following TLR stimulation than those from LPS unexposed mice, characteristic of a trained phenotype. Cytokine production *ex vivo* correlated with atherosclerosis severity *in vivo*. Prenatal LPS did not affect cytokine production capacity. Combined prenatal and postnatal LPS exposure was associated with a reduction in common myeloid progenitor cells in the bone marrow.

**Conclusions:** Postnatal inflammation results in a trained phenotype in atherosclerosis-prone mice that is not enhanced by prenatal inflammation. If analogous mechanisms occur in humans, then there may be novel early life opportunities to reduce CVD risk in infants with early life infections.

## Introduction

Atherosclerosis may develop from early childhood onwards, but manifests symptomatically as CVD in adulthood [1, 2] and it remains the most common cause of death globally [3, 4]. Preterm infants are at increased risk for developing CVD in adulthood [5, 6], with some estimates suggesting a doubling of risk compared to those born at term [7]. This heightened risk may reflect the increased inflammatory exposures that characterise pregnancies that culminate in preterm birth, including prenatal inflammation (chorioamnionitis) and that arising from early life infections, which are highly prevalent in preterm infants [8]. We have previously reported that experimental prenatal and postnatal inflammation both result in additive, deleterious effects on the development of atherosclerosis in *Apoe*^−/−^ mice [9], but the underlying immunological mechanisms are unknown.

Trained immunity, a heightened innate immune response upon exposure to a secondary immunological stimulus as a consequence of epigenetic and metabolic reprogramming, has been implicated in mediating cardiovascular risk [10]. A trained phenotype has been observed in primary human monocytes following exposure to atherogenic stimuli, resulting in increased responsiveness (pro-atherogenic cytokine production) and capacity for foam cell formation and development [11]. In patients with atherosclerosis, monocytes also exhibit a trained phenotype compared to healthy controls, with increased cytokine production and metabolic and epigenetic changes [12]. As cells of the innate immune system only live hours-to-days and the trained phenotype can last up to a year [13], attention has now shifted to their progenitor cells in the bone marrow. Bone marrow progenitor cells are increasingly recognized for their capacity to develop a trained phenotype [14–16], subsequently resulting in changes in peripheral cells. Reprogramming of bone marrow progenitor cells was recently shown in adult patients with atherosclerosis [17]. Risk factors for atherosclerosis, including obesity and diabetes, are associated with subclinical chronic exposure to lipopolysaccharide (LPS), which is a strong inducer of trained immunity [18–23]. To date however, trained immunity and bone marrow reprogramming have only been studied in adults. We hypothesize that early life LPS exposure can induce trained immunity and potentiate atherosclerosis. This may contribute to the association between preterm birth and increased risk of CVD, due to higher incidence of early life inflammation/infection.

Here, we investigate if prenatal and postnatal LPS exposure (experimental surrogates of chorioamnionitis and recurrent childhood infection, respectively) induce a trained phenotype in in peripheral cells and bone marrow progenitors in *Apoe*^−/−^ mice.

## Materials and Methods

### Mice

Animal work was approved by the Monash Medical Centre (MMC) Animal Ethics Committee A (MMCA/2011/36 and MMCA/2011/37). *Apoe*^−/−^ mice were housed in the MMC Specific Pathogen Free animal facility and maintained under alternating 12-hour light/dark periods. Induction of prenatal and postnatal inflammation in these mice has been described previously [9]. In brief, prenatal *Apoe*^−/−^ mice (E15.5) were given intra-amniotic injections (i.a) of either saline or LPS (1μg in 5μL saline), which causes placental inflammation [9]. After birth (E18.5) and 4 weeks of weaning, mice from each group were re-stimulated with LPS (InvivoGen, USA) or PBS (Gibco, USA) at 8, 9, 10, 11, and 12 weeks (5μg in 50μL saline) to mimic recurring childhood infection seen following preterm birth. At 13 weeks, mice were killed humanly.

#### Collection of Peritoneal Macrophages

A midline ventral incision was made from the abdomen up to the head of the mouse and the skin was separated from the peritoneum. 5mL PBS (Gibco) was injected into the peritoneal cavity just below the liver, massaged, redrawn to collect macrophages, and transferred to a 15mL falcon tube (Corning, USA). Cells were pelleted by centrifugation and resuspended in 1mL RPMI (Gibco) for counting using Trypan Blue (Gibco) and a Neubauer Improved cell counting chamber (Sigma-Aldrich, USA). Samples were re-pelleted, resuspended in 2mL freezing medium (90% FCS (Gibco) and 10% DMSO (ATCC, USA)) and transferred to 2mL cryovials for freezing in a Mr Frosty™ Freezing container (Thermo Fisher Scientific, USA) at −80°C.

#### Collection of Bone Marrow Cells

whole bone marrow was extracted from the femurs and tibias of the mice. Feet and muscles were removed, and femur and tibia separated. The tips of each bone were broken using toothed forceps. A single cell suspension was made by flushing the bone marrow out with PBS (Gibco) using a 27-gauge needle and 5mL syringe. Cells were collected onto a petri dish (Thermo Fisher Scientific), transferred to a 50mL Falcon tube (Corning) and pelleted by centrifugation for 10 minutes at 1200rpm. Supernatant was discarded and cells resuspended in 3mL RBC lysis buffer (40.1g NH_2_Cl, 4.2g NaHCO_3_, and 1.8g EDTA in 500mL MilliQ water). Following one minute of incubation, 13mL of RPMI (Gibco) was added and cells re-pelleted. Live cell count was determined using trypan blue, and cells were cryopreserved as described above.

#### Differentiation of bone marrow into bone marrow-derived macrophages

Ten million cells were differentiated into bone marrow-derived macrophages for stimulation. The bone barrow was seeded in 10mL of growth media (GlutamaX 2mM (Gibco), 1% Pen-Strep (Gibco) 10% FCS (Gibco) in RPMI) and 50ng/mL CSF-1 (STEMCELL Technologies, Canada) in petri dishes (Thermo Fisher Scientific). Four days later, 10mL of fresh media was added and on day six after seeding cells were harvested. Cells were scraped in 2mL of cold RPMI (Gibco) and the dish washed with 1mL cold PBS (Gibco). Cells were washed and rested for 2-4 hours prior to stimulation.

#### Stimulations with Pam3Cys, LPS and RPMI

Prior to stimulation or flow cytometric staining, cells from each vial were thawed in a water bath at 37°C and transferred to 10 mL thawing medium 10% FCS (Gibco) 1% Pen-Strep (Gibco) RPMI+Glutamax (Gibco). Live cell count was determined using Trypan Blue (Gibco) and a Countess™ cell counter (Life Technologies, USA). For peritoneal macrophages and bone marrow-derived macrophages, 50,000 cells and 100,000 cells/well were aliquoted for stimulations, respectively. Cells were stimulated in duplicate with LPS (10ng/mL, InvivoGen), Pam3Cys (10μg/ml, InvivoGen), or RPMI (control, Gibco), for 24 hours at 37°C 5% CO_2_. Supernatant from each well was transferred to fresh micronics (Eppendorf, Germany) for storage at −80°C.

#### Assessing Cytokine Concentration

Cytokine levels in supernatants from stimulated cells were determined using the LEGENDplex™ Mouse anti-viral response panel (Biolegend, USA) as per manufacturer’s instructions and read on BD Accuri™ C6 Plus Flow Cytometer (BD Biosciences). Data were analysed using the provided LEGENDplex™ Software (Biolegend).

#### Flow Cytometric analysis of bone marrow myeloid progenitor subsets

Thawed bone marrow was resuspended in 1mL RBC lysis buffer and incubated for 1 minute at 25°C. PBS (25mL, Gibco) was added to suspension and centrifuged prior to resuspension in thawing medium. Five million cells were plated on a round-bottom 96-well plate. Samples were centrifuged at 400 rcf for 5 minutes and incubated for 20 minutes in 30uL 0.2μg/uL (6μg) of CD16/32 PerCP-Cy5.5 (BD Biosciences, USA). Samples were washed and then incubated in 30μL of a 6-colour panel and Ms CD16/32 FACS Block (see Table S1). The gating strategy is shown in Figure 5A. Single-colour compensation controls were either prepared using remaining bone marrow (500,000 cells/well) or using rat/hamster compensation beads (BD anti-rat, anti-hamster Ig κ/negative control set, BD Biosciences). Both compensation controls and samples were washed twice with 200μL FACS buffer (2% FCS (Gibco) in PBS (Gibco)), then resuspended in 200μL FACS buffer and transferred to 5mL FACS tubes for analysis on a BD FACSAria Fusion (SORP) at the Translational Research Institute (Brisbane, Australia). Percentages of hematopoietic stem cells (HSPCs), common myeloid progenitors (CMPs) and granulocyte-macrophage progenitors (GMPs) were quantified (Figure 5A).

#### Statistical Analysis

Statistical analyses were performed using GraphPad Prism, version 8.0.1 for Macintosh (GraphPad Software, La Jolla, CA, USA). Outliers were determined using the ROUT test (p≤0.05). Normal distribution was assessed by the Shapiro-Wilk test. For three or more groups a One-way ANOVA was used if data points were normally distributed, or Kruskal-Wallis test of samples with multiple comparison if data points were not. For assessment of correlations between variables, R values and significance was tested by Pearson’s correlation coefficients if data points were normally distributed, or Spearman’s correlation coefficients if data points were not. A two-sided *p*-value below 0.05 was considered statistically significant.

## Results

### Postnatal LPS treatment induces immune training in peritoneal macrophages which correlates with atherosclerosis

Peritoneal macrophages from mice treated postnatally with LPS were stimulated *ex vivo* with Pam3Cys or LPS. Following 24-hour stimulations with toll-like receptor (TLR) 1/2 agonist Pam3Cys, peritoneal macrophages from mice treated postnatally with LPS *in vivo* exhibited significantly higher cytokine responses compared to peritoneal macrophages derived from saline-treated controls (Figure 1A). This cytokine response was also strongly associated with CD45^+^ cell infiltration into atherosclerotic plaques (Figure 1B). These same trends were observed upon *ex vivo* stimulation with LPS (Figure S2).

**Figure 1.**
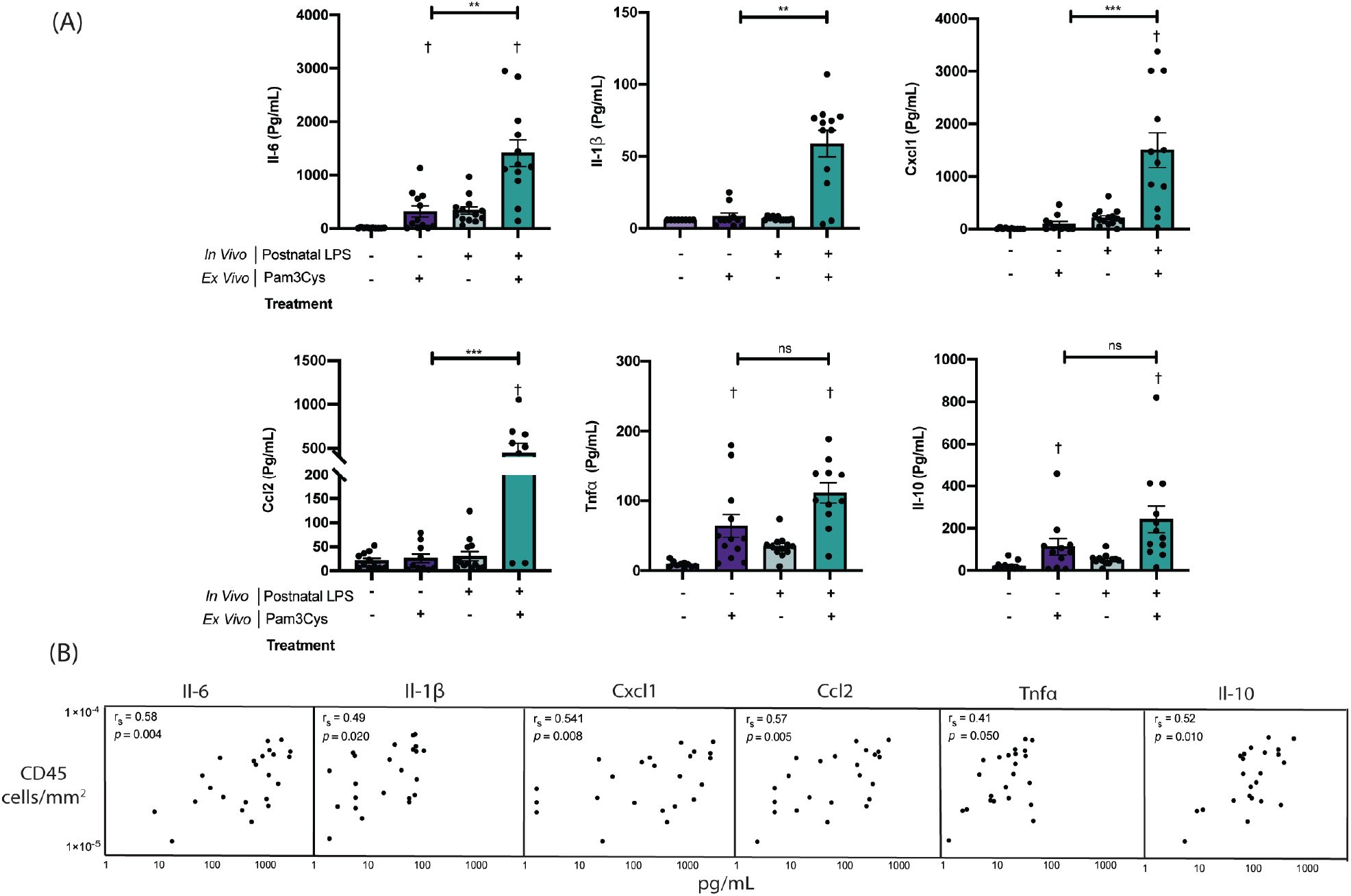
Postnatal LPS in Apoe^−/−^ mice results in increased cytokine response of peritoneal macrophages following *ex vivo* stimulation with Pam3Cys. **(A)** Cytokines recorded in supernatant following stimulation of peritoneal macrophages with Pam3Cys (10ug/mL) or RPMI for 24 hours at 37°C 5% CO_2_. Data represents mean ± SEM with each point representing results from one mouse. Outliers removed using ROUTS test (*p*<0.05). For all panels, statistical significance was tested using a student’s t-test or Mann-Whitney test as appropriate. Statistical significance between treatment groups is denoted by * (*p*<0.05) and *** (*p*<0.001). Statistical significance between cells stimulated *ex vivo* with Pam3Cys and those stimulated with RPMI (from the same mice) is denoted by † (*p*<0.05). **(B)** Table presents correlation between CD45^+^ cell infiltration (cells/m^2^) into atherosclerotic plaque and cytokine response of peritoneal macrophages (Pam3Cys) from mice postnatally treated with saline or LPS. R values and significance tested by Pearson’s correlation coefficient.

### Postnatal LPS treatment induces immune training in bone marrow-derived macrophages

The observed hyperinflammatory phenotype may be the result of immune training in the myeloid progenitors of these cells located in the bone marrow [14]. To examine this possibility, bone marrow-derived macrophages were derived *ex vivo* and stimulated with TLR4 agonist LPS. Following 24-hour stimulations, bone marrow-derived macrophages from mice treated postnatally with LPS show a trend of higher cytokine responses compared to those derived from saline-treated controls. However, this was only statistically significant for Gm-csf, and was not associated with increased CD45^+^ cell infiltration into atherosclerotic plaques in these mice (Figure 2).

**Figure 2.**
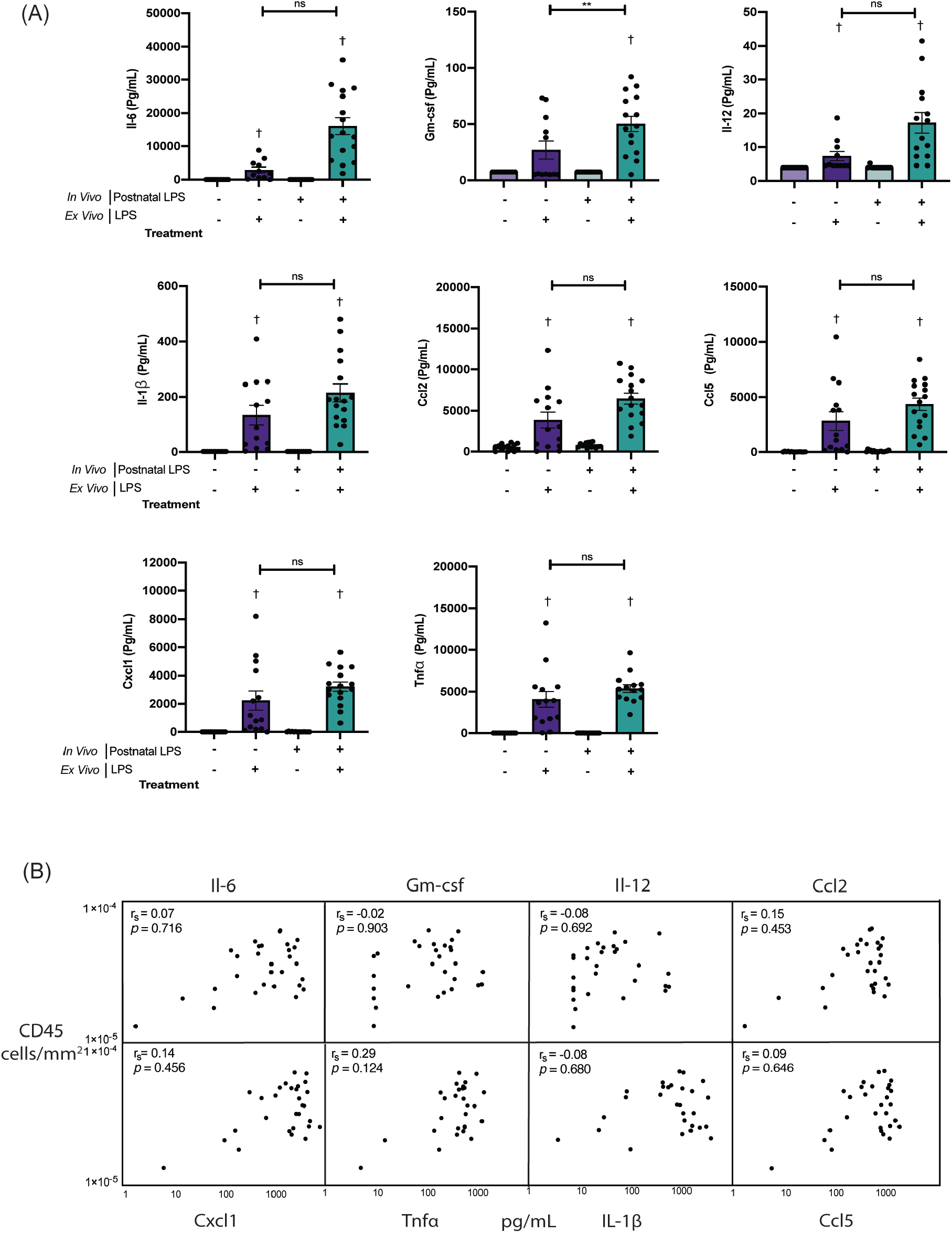
Postnatal LPS in *Apoe*^−/−^ mice results in increased Gm-csf response of bone marrow-derived macrophages following *ex vivo* stimulation with LPS. **(A)** Cytokines recorded in supernatant following stimulation of peritoneal macrophages with LPS (10ng/mL) or RPMI for 24 hours at 37°C 5% CO_2_. Data represents mean ± SEM with each point representing results from one mouse. Outliers removed using ROUTS test (*p*<0.05). For all panels, statistical significance was tested using a student’s t-test or Mann-Whitney test as appropriate. Statistical significance between treatment groups is denoted by * (*p*<0.05) and *** (*p*<0.001). Statistical significance between cells stimulated *ex vivo* with LPS and those stimulated with RPMI (from the same mice) is denoted by † (*p*<0.05). **(B)** Table presents correlation between CD45^+^ cell infiltration (cells/m^2^) into atherosclerotic plaque and cytokine response of bone marrow-derived macrophages (LPS) from mice postnatally treated with saline or LPS. R values and significance tested by Pearson’s or Spearman’s correlation coefficient as appropriate.

### Prenatal LPS treatment does not affect the immune training induced by postnatal LPS in peritoneal macrophages

Given that the combination of prenatal and postnatal inflammation induced greater atherosclerosis compared to either treatment alone [24], we next sought to determine if this was due to heightened immune training compared to either treatment alone. Peritoneal macrophages derived from mice treated only prenatally with LPS exhibited no increase in cytokine expression relative to controls following 24-hour *ex vivo* stimulation with Pam3Cys (Figure 3). Importantly, whilst peritoneal macrophages from mice treated *both* pre- and postnatally with LPS exhibited significantly higher cytokine responses to all stimuli when compared to saline treated controls, these levels were not different from mice treated *only* postnatally (Figure 3). These same results were observed upon *ex vivo* stimulation with LPS (Figure S2).

**Figure 3.**
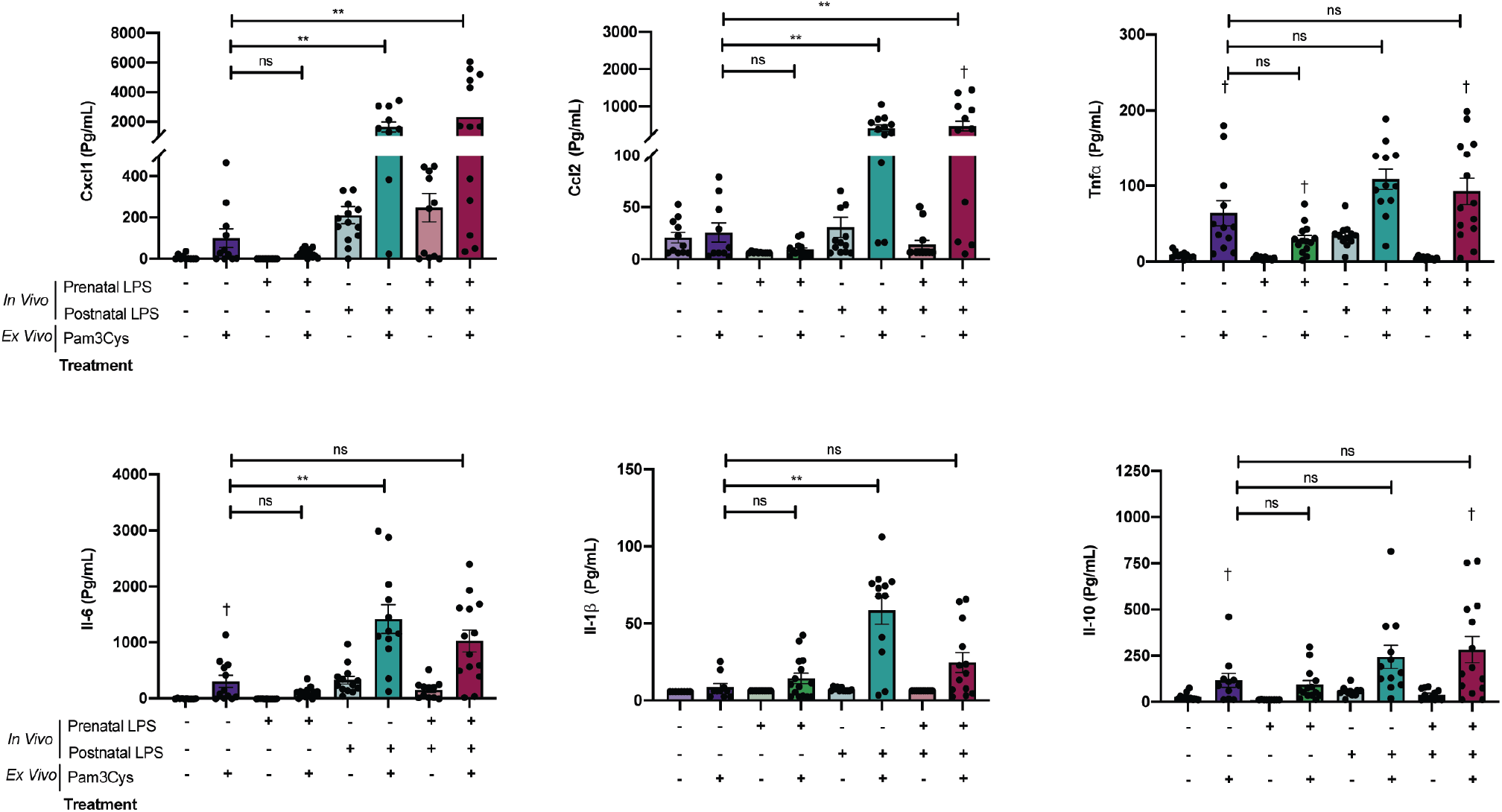
Prenatal LPS in *Apoe*^−/−^ mice does not result in increased cytokine response of peritoneal macrophages following *ex vivo* stimulation with Pam3Cys. **(A)** Cytokines recorded in supernatant following stimulation of peritoneal macrophages with Pam3Cys (10ug/mL) or RPMI for 24 hours at 37°C 5% CO_2_. Data represents mean ± SEM with each point representing results from one mouse. Outliers removed using ROUTS test (*p*<0.05). Statistical significance was tested using a student’s t-test or Mann-Whitney test as appropriate. Statistical significance between treatment groups is denoted by * (*p*<0.05) and ** (*p*<0.01). Statistical significance between cells stimulated *ex vivo* with Pam3Cys and those stimulated with RPMI (same mouse) is denoted by † (*p*<0.05).

### Prenatal LPS treatment does not affect the immune training by postnatal LPS in bone marrow-derived macrophages

We next studied the effect of prenatal LPS on the immune response of bone marrow-derived macrophages and their progenitors. Bone marrow-derived macrophages from mice treated only prenatally with LPS exhibited no increase in cytokine response relative to controls following 24-hour *ex vivo* stimulation with LPS (Figure 4). There was also no additive effect on cytokine response when prenatal LPS was followed by postnatal treatment with LPS (Figure 4). These data suggest that prenatal exposure to LPS has minimal effect on the observed pro-inflammatory response in peritoneal macrophages or bone marrow-derived macrophages. Another training-related mechanism may explain the additive effect of prenatal LPS exposure on the atherosclerosis phenotype [9].

**Figure 4.**
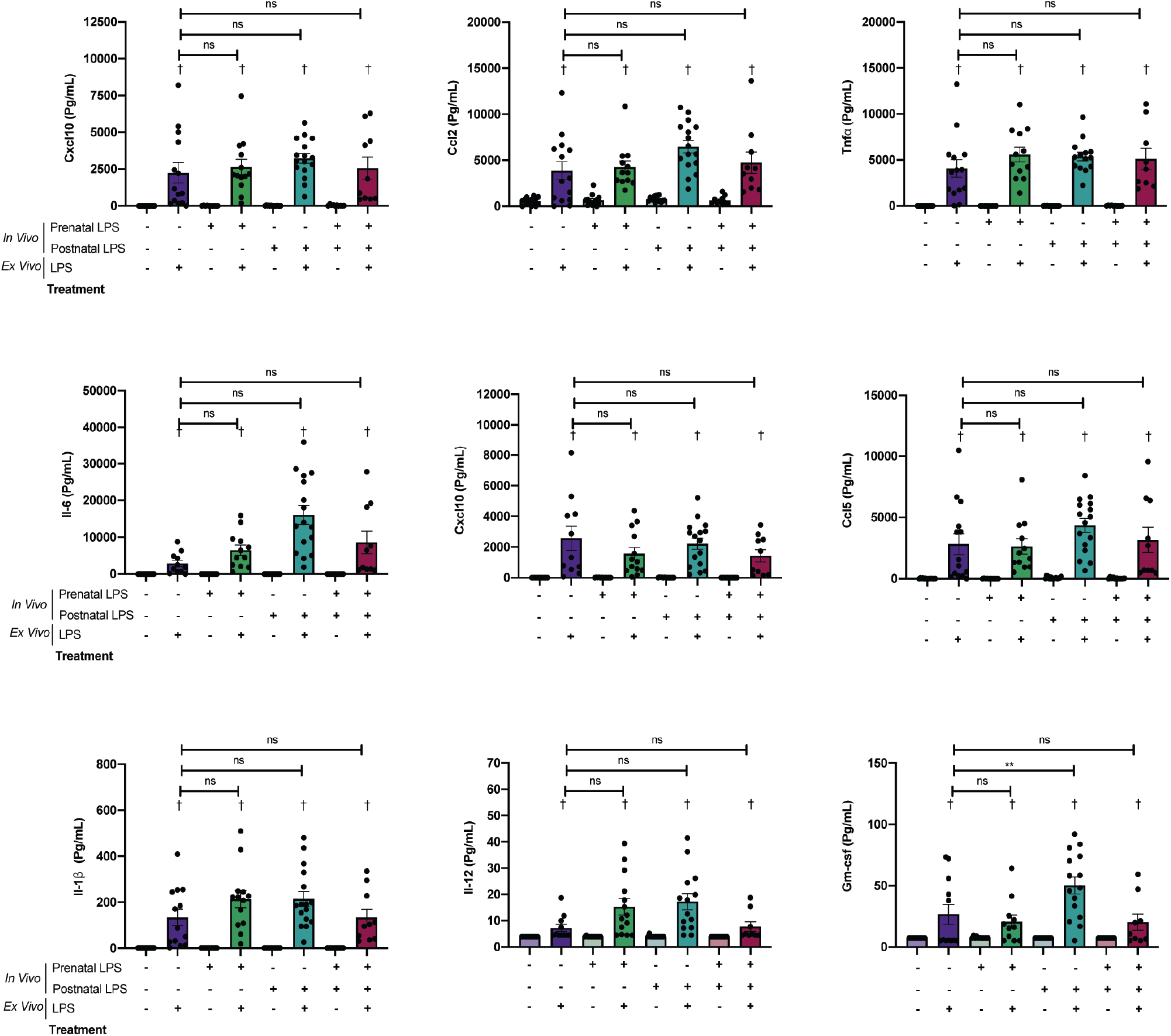
Prenatal LPS in Apoe^−/−^ mice does not result in increased cytokine response of bone-marrow derived macrophages following *ex vivo* stimulation with LPS. Cytokines recorded in supernatant following stimulation of bone marrow-derived macrophages with LPS (10ng/mL) or RPMI for 24 hours at 37°C 5% CO_2_. Data represents mean ± SEM with each point representing results from one mouse. Outliers removed using ROUTS test (*p*<0.05). Statistical significance was tested using a student’s t-test or Mann-Whitney test as appropriate. Statistical significance between treatment groups is denoted by * (*p*<0.05) and ** (*p*<0.01). Statistical significance between cells stimulated *ex vivo* with LPS and those stimulated with RPMI (same mouse) is denoted by † (*p*<0.05).

### Prenatal and postnatal treatment alters bone marrow progenitor populations

We sought to determine if the combination of prenatal and postnatal inflammation facilitated atherosclerosis via another mechanism rather than enhanced immune training. It was reasoned that changes in bone marrow populations could result in alterations in numbers or functioning of circulating monocytes which could contribute to the atherosclerosis phenotype observed [14]. Whilst prenatal and postnatal LPS treatment alone did not produce any significant change in progenitor populations relative to untreated mice, prenatal and postnatal LPS treatment together significantly reduced progenitor cell subset populations (CMPs and GMPs) (Figure 5B).

**Figure 5.**
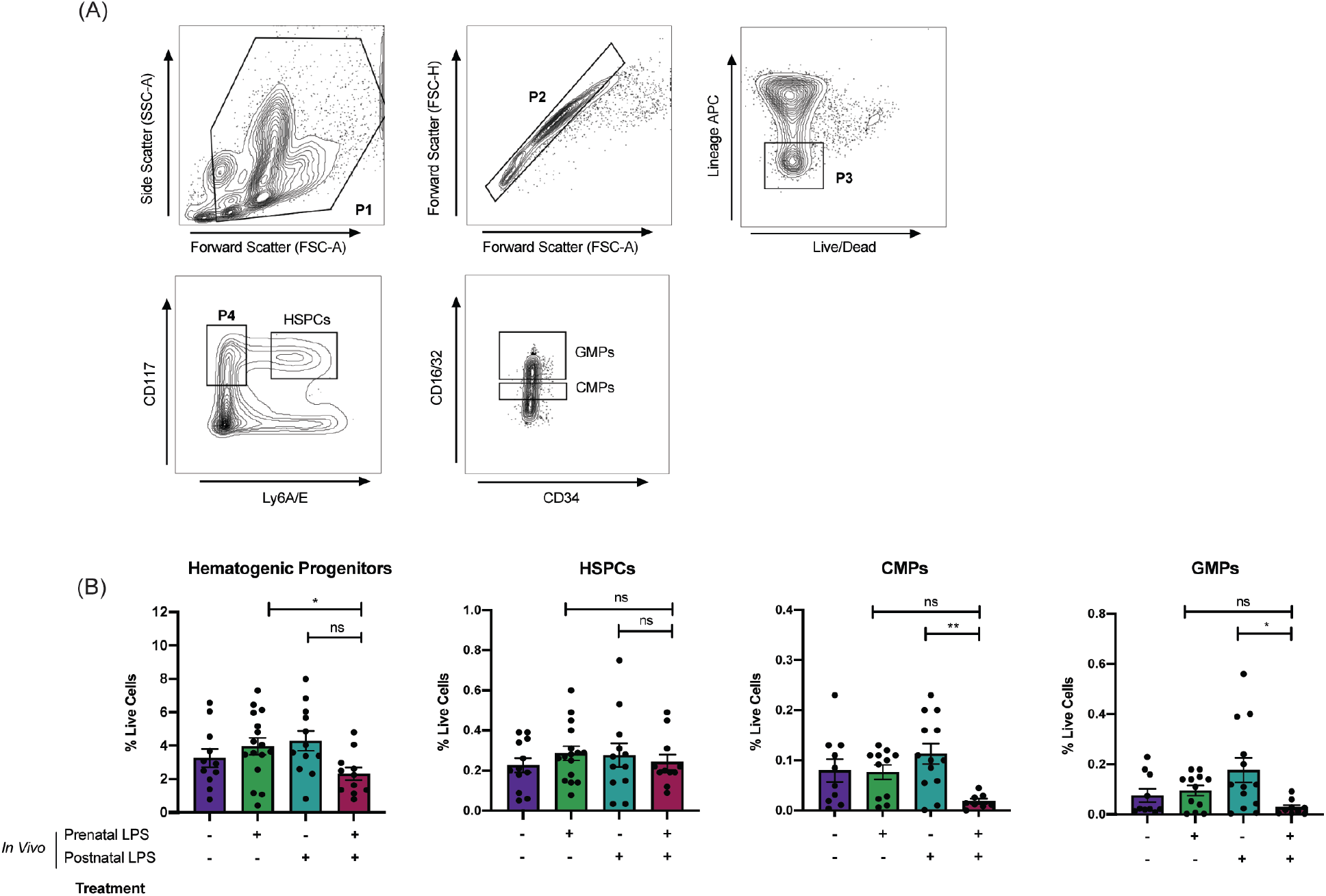
Combined Pre- and Postnatal treatment of LPS in Apoe^−/−^ mice results in a reduction of CMP myeloid progenitor populations in the bone marrow. **(A)** Gating strategy for bone marrow HSPC, CMP and GMP populations. Leucocytes were separated using forward/sideward scatter. Live/progenitor cells were negatively selected. We distinguished HSPCs (Sca-1+ c-kit^+^), GMPs (CD34^int^, CD16/32^high^) and CMPs (CD34^int^, CD16/32^int^). **(B)** Frequency of total hematogenic progenitors, HSPC, CMP, and GMP populations in bone marrow as a percentage of live cells. Data represents mean ± SEM with each point representing results from one mouse. Outliers were removed using ROUTS test (*p*<0.05). For all panels, statistical significance between all groups and untreated mice was tested using an analysis of variance (ANOVA) or Kruskal-Wallis test as appropriate and is denoted by * (*p*<0.05) and ** (*p*<0.01).

## Discussion

Both prenatal and repeated postnatal inflammation have deleterious and additive effects on atherosclerosis development in *Apoe*^*-/*-^ mice, characterised by increased atherosclerotic lesion number, size and severity [9]. This phenotype was independent of plasma lipid levels [9], suggesting that immune mechanisms likely underlie the observed changes. Here, we provide evidence that in *Apoe*^*-/*-^ mice postnatal LPS stimulation induces a trained phenotype *ex vivo* and that this is associated with increased severity of atherosclerosis plaques. We report for the first time that prenatal LPS stimulation does not induce a trained phenotype *ex vivo*, but the combination of prenatal and postnatal inflammation does result in a reduction of BM progenitor subset populations.

The role of inflammation in CVD is widely recognised and inflammatory pathways are rapidly emerging as attractive therapeutic targets [25, 26]. Notably, postnatal inflammation (from both infectious and non-infectious stimuli) is associated with atherosclerosis later in life [27]. Trained immunity has been proposed as a mechanism underlying these epidemiological associations [28, 29]. Here, we demonstrated that recurring postnatal inflammation induced a trained phenotype in peritoneal macrophages of *Apoe*^−/−^ mice, and this hyperinflammatory response was associated with increased severity of atherosclerosis. These results corroborate previous findings, where *Apoe*^−/−^ mice fed a high-fat diet and treated with subclinical LPS developed polarised, pro-inflammatory monocytes, which subsequently accelerate atherosclerosis [30]. Importantly, these results are also consistent with findings from human studies in adults. Monocytes from patients with elevated LDL-cholesterol have increased secretion of pro-inflammatory cytokines in response to ex-vivo stimulation with TLR antagonists [31], a phenotype that did not change upon lowering of LDL-cholesterol (i.e. trained phenotype). Moreover, a trained phenotype can be induced in human monocytes following exposure to atherogenic stimuli (e.g. oxLDL), which then contributes to atherosclerosis through increased foam cell formation and proinflammatory cytokine production [11].

As circulating monocytes have a short lifespan [32], it has been suggested that training begins in the bone marrow and is conferred to daughter cells [14–16]. In this study, the trained phenotype observed in peritoneal macrophages following *ex vivo* stimulation was weakly present in bone marrow-derived macrophages, although not associated with severity of atherosclerosis. Consequently, further studies confirming the central (i.e. bone marrow) origins of this phenotype following postnatal inflammation, and impacts on atherogenesis, are warranted.

Prenatal inflammation (i.e. chorioamnionitis) has also been implicated in development of atherosclerosis [33, 34], but links to trained immunity have been inconsistent. For example, in humans prenatal inflammation alters the neonatal transcriptome with activation of innate immune pathways [35], consistent with studies in rats [36]. However, a hyporesponsive transcriptional profile in cord blood monocytes of preterm infants has also been observed, coupled with decreased pro-inflammatory cytokine expression upon secondary stimulation [37]. The current study suggests that in this *Apoe*^−/−^ model, prenatal inflammation *per se* does not result in induction of a trained phenotype in peritoneal macrophages or bone marrow derived macrophages, and suggests an alternative explanation for the exacerbated atherosclerosis previously observed in mice with both pre- and postnatal inflammation [24]. It should be noted, however, that a similar study in rats found PBMCs exhibited a proinflammatory phenotype at P21 following birth [36], so the possibility that a trained phenotype could be observed at a different timepoint following postnatal LPS exposure should not be excluded.

Interestingly, the combination of pre- and postnatal LPS resulted in reduction in percentages of common myeloid progenitors in the bone marrow, compared to mice only treated postnatally. This is unlike that observed in previous models of trained immunity, which typically feature increased frequency of myeloid progenitor subsets [16]. Reduction in these populations could be indicative of heightened myelopoiesis of these populations (which have limited self-renewal capacity) [38], or the migration of myeloid progenitors (HSPCs) to the spleen where they undergo enhanced extramedullary myelopoiesis. Either could result in increased circulating monocytes, which is supported by the significant increase in CD45^+^ leukocyte infiltration in plaques in these mice [24].

The present study provides experimental evidence of trained immunity as a mechanistic link between early-life postnatal inflammation and development of CVD [14]. However, it is important to recognize limitations of the *Apoe*^−/−^murine model and its relevance to human disease remains debated [39, 40]. In addition, although *Apoe*^−/−^ mice are a well-established transgenic model of atherosclerosis, the *Apoe* gene is expressed on several immune cells including macrophages and has anti-inflammatory properties [41]. Finally, differences with the methodology of this study and the pathogenesis of prenatal/postnatal inflammation should also be acknowledged; chorioamnionitis and postnatal inflammation following preterm birth does not result simply from LPS exposure, but rather from a variety of pathogens and PAMPS [42–45].

Despite these limitations, potential clinical implications of these data should be considered. The population of ex-preterm infants is increasing, and they are at increased CVD risk. Chorioamnionitis is implicated in 40-70% of preterm births [46], and preterm individuals are at significantly higher risk for childhood infection [47]. If analogous mechanisms are present in human preterm infants, these data suggest that children with a history of repeated early life inflammation may be at risk of accelerated atherosclerosis. Human studies to assess the role of early life inflammation, immune training and the subsequent development of CVD are needed.

## Author Contributions

E.C.N. co-wrote manuscript, performed experimental protocols and analysis

S.B, D.P B., K.R.S. designed the research, performed data analysis and co-wrote the manuscript

T.J.M. contributed to methodology and design of research

H. L., M.J.W., A.P. L., M.U.N., L.K.W. S.P., M.M.C., T.R.M. contributed to experimental work and data analysis

Funding acquisition: T.J. Moss and D.P B.

## Supplementary Materials

**Table S1.** Cell markers and their respective antibodies for investigating bone marrow progenitor populations of interest in *Apoe*^−/−^ mice.

| Marker | Antibody | BD Biosciences Product Number | Clone | Concentration Used |
| --- | --- | --- | --- | --- |
| CD117 (C-kit) | Ms CD117<br>BUV395 2B8 | 564011 | 2B8 | 1:400 |
| CD34 | Ms CD34 PE<br>RAM34 | 551387 | RAM34 | 1:50 |
| Ms CD16/32 | Ms CD16/32<br>PerCp-Cy5.5 2.4 G2 | 560540 | 2.4G2 | 1:200 |
| Ly6A/E (Sca-1) | Ms Ly6A/E<br>BV421 D7 | 562729 | D7 | 1:400 |
| Lineage Cocktail | Ms Lineage Cocktail APC with isotype control | 558074 | RA3<br>TER-119<br>145-2C11<br>M1/70<br>RB6-8C5 | 1uL/sample |
| Live/Dead | Fixable Viability Stain 700 | 564997 |  | 1:15000 |

**Figure S1.**
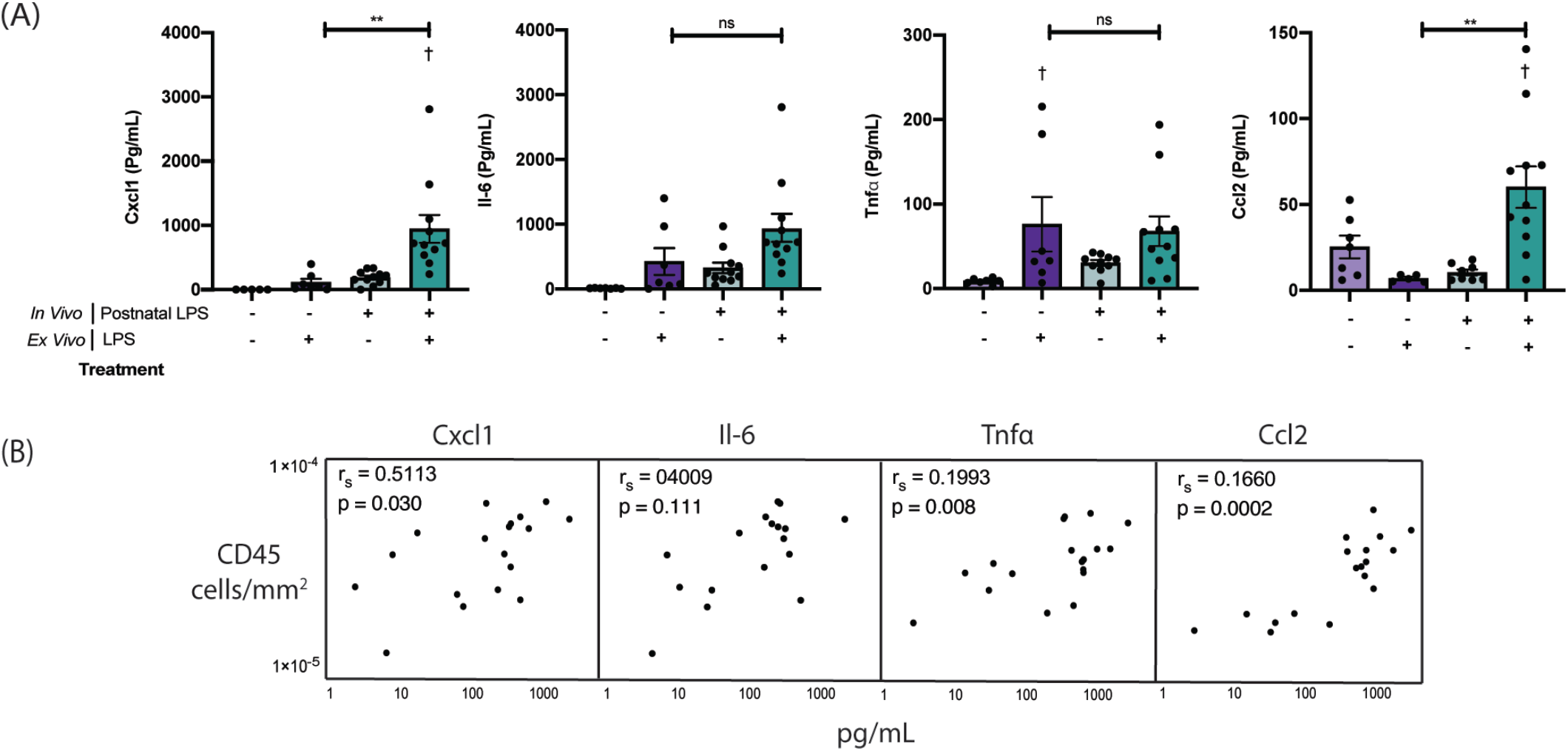
Postnatal LPS in Apoe^−/−^ mice results in increased cytokine response of peritoneal macrophages following *ex vivo* stimulation with LPS. **(A)** Cytokines recorded in supernatant following stimulation of PMs with LPS (10ng/mL) or RPMI for 24 hours at 37°C 5% CO_2_. Data represents mean ± SEM with each point representing results from one mouse. Outliers removed using ROUTS test (*p*<0.05). For all panels, statistical significance was tested using a student’s t-test or Mann-Whitney test as appropriate. Statistical significance between treatment groups is denoted by * (*p*<0.05) and *** (*p*<0.001). Statistical significance between cells stimulated *ex vivo* with LPS and those stimulated with RPMI (from the same mice) is denoted by † (*p*<0.05). **(B)** Table presents correlation between CD45^+^ cell infiltration (cells/m^2^) into atherosclerotic plaque and cytokine response of PMs (LPS) from mice postnatally treated with saline or LPS. R values and significance tested by Pearson’s correlation coefficient.

**Figure S2.**
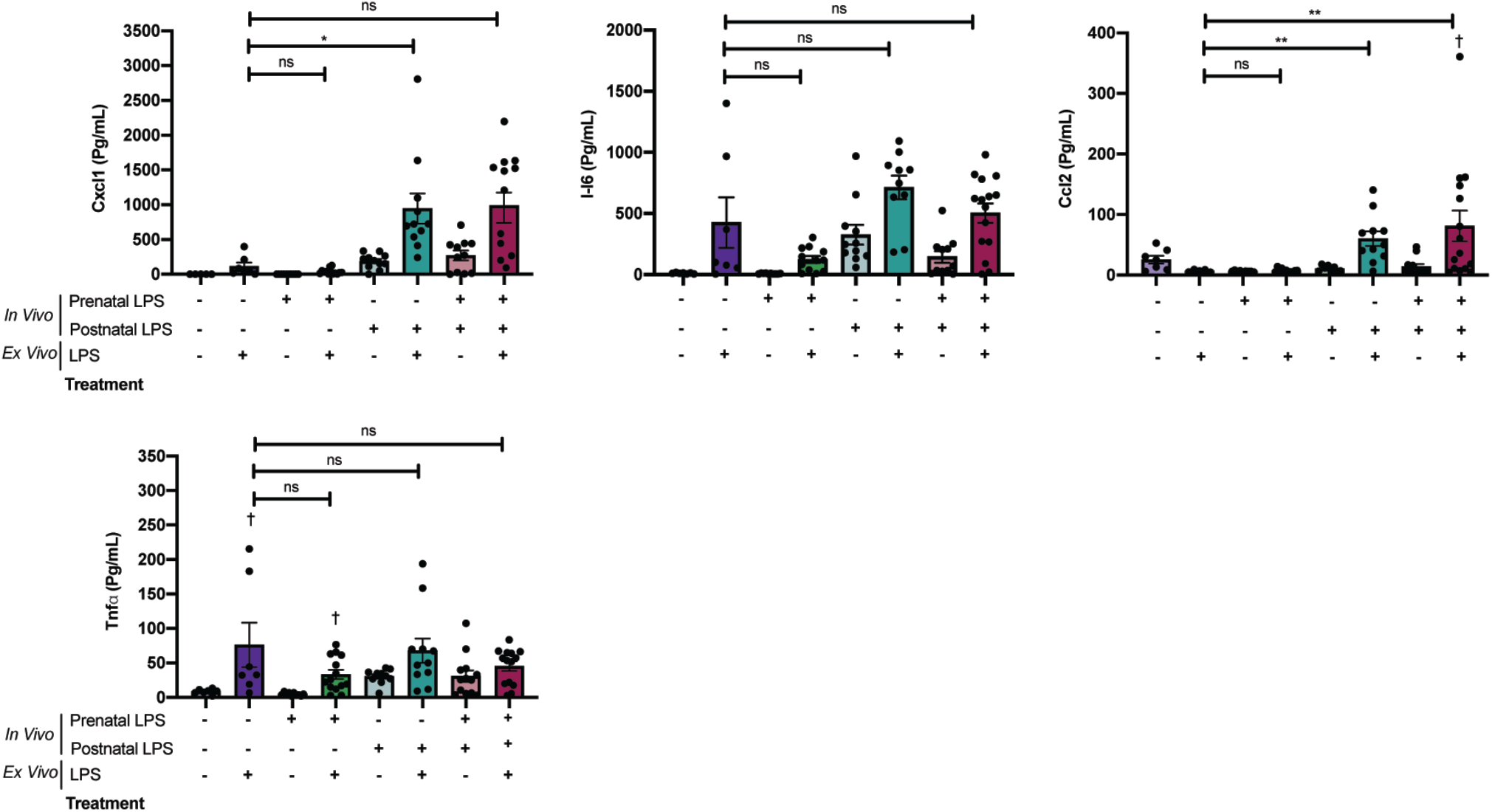
Postnatal LPS in Apoe^−/−^ mice results in increased cytokine response of peritoneal macrophages following *ex vivo* stimulation with LPS. Cytokine response is associated with atherosclerosis. **(A)** Cytokines recorded in supernatant following stimulation of PMs with LPS (10ng/mL) or RPMI for 24 hours at 37°C 5% CO_2_. Data represents mean ± SEM with each point representing results from one mouse. Outliers removed using ROUTS test (*p*<0.05). Statistical significance was tested using a student’s t-test or Mann-Whitney test as appropriate. Statistical significance between treatment groups is denoted by * (*p*<0.05) and ** (*p*<0.01). Statistical significance between cells stimulated *ex vivo* with LPS and those stimulated with RPMI (same mouse) is denoted by † (*p*<0.05).

